# NN-Assisted Image Analysis for Quantifying Intracellular *Trypanosoma cruzi* Infection

**DOI:** 10.64898/2026.02.28.707609

**Authors:** Joaquín Iolster, Salomé Catalina Vilchez Larrea, Guillermo Daniel Alonso

## Abstract

Quantification of intracellular *Trypanosoma cruzi* infection remains a central, yet methodologically challenging step in Chagas disease research and early-stage drug discovery. Current approaches largely rely on manual microscopy-based counting or on genetically modified parasites, both of which present limitations in scalability, reproducibility, or accessibility. Here, we developed and validated a neural network (NN)-based pipeline for the automated quantification of infection rates and parasite burden in mammalian cells using images stained exclusively with DNA-binding fluorescent dyes. Two independently refined deep-learning models were trained to segment host cell nuclei and intracellular amastigotes, respectively, and subsequently integrated into a unified algorithm that assigns each parasite to its nearest host cell. The pipeline was evaluated using confocal images from six mammalian cell lines infected with two *T. cruzi* strains and compared against blinded manual quantification. Automated detection of both, host nuclei and parasites, showed high concordance with manual counts, with median deviations around 5% and similar distributions of parasite burden per cell. In contrast to morphology-based image analysis methods, our NN-based approach demonstrated improved robustness across diverse cell types and staining conditions, reduced parameter dependency, and independent segmentation of host and parasite objects, minimizing error propagation. Although minor biases in parasite-to-cell assignment were observed in densely clustered cultures, overall infection indexes and burden estimates closely matched manual analysis. This accessible and scalable AI-assisted workflow provides a reproducible alternative to manual quantification and represents a methodological advance for standardized phenotypic screening of intracellular *T. cruzi*, supporting more robust and harmonized drug discovery efforts in Chagas disease.

## Introduction

Chagas disease, caused by the protozoan parasite *Trypanosoma cruzi*, remains one of the most prevalent neglected tropical diseases in Latin America, with increasing global relevance due to migration patterns (1). Benznidazole (BNZ) and nifurtimox (NFX), the two drugs currently approved for the treatment of *T. cruzi* infection, exhibit trypanocidal activity and are effective during the acute phase of the disease. However, their efficacy is inconsistent in the chronic phase, and they fail to improve clinical outcomes once irreversible organ damage has occurred (2). The limited efficacy observed during the chronic phase is likely attributable to the presence of parasite forms with reduced replication rates and increased resistance to current therapies (3). In addition, the relatively high frequency of adverse effects associated with existing treatments, together with the lack of a vaccine, underscores the urgent need for novel therapeutic strategies which must be assessed through well-established methodologies ensuring robust reproducibility (2).

*T. cruzi* exhibits a complex life cycle that includes a crucial intracellular stage within nucleated host cells. During this stage, trypomastigotes invade host cells, differentiate into amastigotes, and proliferate within the cytoplasm before transforming back into trypomastigotes and lysing the cell (4). The intracellular amastigote stage is of paramount importance for the progression of the disease and constitutes the primary target for antiparasitic drug development.

The initial step in the drug discovery process involves the identification of compounds with potent and selective antiparasitic activity through in vitro phenotypic screening cascades (5). Despite its biological and therapeutic relevance, evaluating the efficacy of drug candidates against the intracellular form of *T. cruzi* remains a major challenge. Genetically modified parasites expressing beta-galactosidase or Firefly luciferase have been widely used in colorimetric or luminometric assays to evaluate *in vitro T. cruzi* infection. This method is simple, quick and inexpensive, rendering it useful for high-throughput screening of large compound libraries to identify new drugs (6). However, the information that can be obtained from this type of assay is limited, as only relative infection rates can be evaluated, missing out on valuable details. Other conventional phenotypic screening assays on infected mammalian cells require microscopic quantification of infection rates, parasite burden, and intracellular development. These assays provide more detailed information but are often labor-intensive, time-consuming, and subject to high inter- and intra-operator variability (7). In this regard, a few algorithms for automated infection quantification of microscopic images have been developed by different research groups (7–9). Moon et al. (8) highlighted the application of image-based high-throughput screening for *T. cruzi* and demonstrated the potential of fluorescence-based microscopy coupled with automated quantification to streamline the assessment of antiparasitic activity. Also in 2014, Yasdanparast et al. (7) implemented an image-based high-content screening strategy for anti-trypanosomal compounds (INsPECT), further validating the utility of automated image analysis for robust and scalable drug screening. The inherent complexity of distinguishing infected cells, accurately counting intracellular amastigotes, and analyzing large numbers of samples significantly limits the scalability and reproducibility of such studies. Moreover, these methods are highly sensitive to the quality of the images and the analysis parameters must be adjusted according to the acquisition parameters.

To overcome these limitations, high-content imaging combined with artificial intelligence (AI)-based image analysis has gained relevance for the quantitative assessment of intracellular infections. Deep learning approaches using convolutional neural networks (CNNs) are particularly powerful, as they can be trained to classify and segment complex cellular structures from microscopy images (10). The inherent complexity of distinguishing infected cells, accurately counting intracellular amastigotes, and analyzing large numbers of samples significantly limits the scalability and reproducibility of such studies. AI-assisted analysis has already been successfully implemented for other parasitic diseases, as well as in cancer and stem cell research (11–14). These technologies have been shown to not only accelerate data acquisition but also enhance reproducibility and enable harmonization of screening protocols across laboratories.

In this work, we present the development of a NN-based algorithm for the quantification of infection rates and parasite burden on microscopy images of *Trypanosoma cruzi*-infected mammalian cells. We trained and tested the ability of two independent NN-based models to detect individual host cells and to accurately distinguish *T. cruzi* parasites on microscopic images solely stained with a DNA-binding dye. We then implemented an algorithm combining both models to assign each parasite to a mammalian cell, in order to retrieve the percentage of infected host cells and the number of parasites per infected cell. The outcome of the algorithm was compared to manual quantification of these parameters in a sub-group of images, showing similar results. We also contrasted the ease-of-use and results of our method to previously published algorithms, which showed our method rendered comparable results. Considering the intracellular amastigote form of *T. cruzi* is the most relevant target for therapeutic intervention, the implementation of automated image analysis tools based on neural networks can represent a significant methodological advance as these approaches support large-scale, reproducible, and high-resolution evaluation of compound activity, aligning with broader efforts to incorporate AI and digital technologies into biomedical research and neglected disease drug development (15,16).

## Material and Methods

### Parasite and mammalian cells

Vero cells (ATCC® CCL-81™), HeLa cells (ATCC® CCL-2™) and Caco2 cells (ATCC® HTB-37™) were cultivated in MEM supplemented with 5% foetal bovine serum (FBS) or 20% FBS (for Caco2 cells), 100 U/ml penicillin, 0.1 mg/mL streptomycin, and 2 mM glutamine. AC16 cells (ATCC® CRL-3568™) and BeWo cells (ATCC® CCL-98™) were cultivated in DMEM-F12 supplemented with 10% or 5% FBS (respectively for AC16 or BeWo cells), 100 U/ml penicillin, 0.1 mg/mL streptomycin, and 2 mM glutamine. THP-1 cells (ATCC® TIB-202™) were cultivated in RPMI supplemented with 10% FBS, 1 mM sodium pyruvate, 0.05 mM 2-mercaptoethanol, 1.4 mM glucose, 100 U/ml penicillin, 0.1 mg/mL streptomycin, and 2 mM glutamine. *T. cruzi* parasites from the Tulahuen and Dm28c strains were maintained in culture by infection of Vero cells in MEM with 5% FBS, 2 mM L-Glutamine and 1% penicillin-streptomycin.

THP-1 cells were differentiated using 80 nM of phorbol 12-myristate 13-acetate (PMA) for 24 hours, after which the PMA was removed and the cells recovered for an additional 48 h.

To obtain the samples for the validation dataset, we seeded coverslips with 2x10^4^ cells. After 24 h, they were infected with trypomastigotes obtained from the supernatants of infected Vero cells at a 40:1 multiplicity of infection. After 4 h, the parasites were removed, the cells thoroughly washed and the corresponding media replaced. We finally fixed the cells at 48 h post infection.

### Fluorescence microscopy

Cells were seeded on glass coverslips in 24 well plates and subjected to the infection scheme. After fixation (PFA 4%, 15 min), nuclei were counterstained with DAPI or Hoechst 33342, and the coverslips were then mounted in Vectashield (Molecular Probes P36930, Eugene, OR, USA). For specific immunostaining, coverslips bearing infected cells were fixed in PFA 4% for 15 min, permeabilized in PBS 1x-Triton X-100 0.1% for 10 min, blocked in PBS 1x-BSA 3% for 1 h and then incubated overnight with anti-*α*-Tubulin monoclonal antibody (1:2000, Sigma-Aldrich T5168) and anti-*T. cruzi* mouse serum (1:1000), kindly gifted by Dr. Karina Gómez. After incubation with the antibody and serum, coverslips were washed three times with PBS 1x - Tween 0.05% (TPBS) and incubated for 2 h with AlexaFluor488 anti-mouse and AlexaFluor546 anti-rabbit antibodies. The samples were then counterstained and mounted as before.

The validation dataset was obtained on a Leica TCS SPE confocal microscope. A 405 nm excitation laser was used for imaging Hoechst 33342, a 488 nm laser for Alexa488 and a 532 nm laser for Alexa546. The detection range for each channel was set using the corresponding emission spectra provided by the manufacturer’s software (Leica LAS X). Images were obtained with a zoom of 1.5 on a 40X/1.1NA objective, at a 1024x1024 pixel resolution.

### Manual quantification of infection rates and parasite burden

The manual quantification of infection rate and amastigotes per cell in the images was done using the multi-point tool of the Fiji software. We started by counting the host cell nuclei. Then, for each individual nucleus, we found the associated parasites and recorded the number. This allowed us to obtain a dataset with the parasite burden for each cell, from which we calculated the summary statistics for each image. To avoid potential biases and assess human error, the names of the images were masked at random so as to hide the treatment they belonged to. The manual quantification of the infection rates and amastigote burden was done by at least two different operators.

### Model training for mammalian cell and *T. cruzi* amastigote detection (Cellpose GUI)

Cellpose was used as the base model for cell segmentation in fluorescence microscopy images (17). Cellpose is a deep-learning based segmentation method designed to precisely segment cells from a wide range of image types with no or minimal parameter adjustment. Using the cellpose GUI, we refined the Cellpose-SAM model with a human-in-the-loop training scheme (18). We used a unique set of training images for either the host cell or the *T. cruzi* model, including images in RGB or grayscale format, obtained by either widefield or confocal microscopy, with magnifications ranging from 20X to 60X. To train the model to detect host-cell nuclei of infected cells, we used 7883 regions of interest (ROIs) from 76 images, while the model for *T. cruzi* detection required 87 images, with 9254 ROIs.

### Algorithm development for quantification of infection rates and parasite burden

The image analysis was automated using the Python programming language. Briefly, the images are loaded and segmented separately with the host nuclei model and the parasite model. The distances between all parasite and host mask centroids are measured and each parasite object is assigned to its nearest host object. The number of parasites in a cell is then recorded in a row of the single-cell dataframe. A separate dataframe also records the summary statistics for all cells in each image. Finally, a plot is generated for every image to later evaluate the quality of the segmentation and assignment.

### Statistical analyses

The number of amastigotes per cell and per infected cell, as well as the proportion of infected cells were modelled with Generalized Linear Mixed Models. Because the first two variables correspond to non-normally distributed count data, we used a negative binomial distribution with a log-link function, and a truncated negative binomial distribution for the number of amastigotes per infected cell. To compensate for the zero-inflation in the amastigotes per cell variable, we set the zi-formula to 1. A similar approach was used for modelling the proportion of infected cells, specifying a binomial distribution (0 = uninfected and 1 = infected), using a logit-link function. All models included the Method (Automatic and Manual), the parasite Strain (Tulahuen and Dm28c) and Cell line (AC16, BeWo, Caco2, HeLa, THP-1 and Vero) as fixed effects variables, along with their three way interaction. The lack of independence between images obtained from the same samples was included as a random intercept.

The models were fitted in R using the glmmTMB package. Simple effects comparisons between the Methods for each Strain-Cell group were carried out with the emmeans package.

### Code availability

All of the custom code and algorithms we have used for these analyses are archived on Zenodo (DOI:10.5281/zenodo.18745819) and available on GitHub (https://github.com/jiolster/NN-Assisted-Image-Analysis-for-Quantifying-Intracellular-Trypanosoma-cruzi-Infection.git).

## Results

### Neural Network training for mammalian cell detection

To simplify sample preparation and increase the method’s applicability, we decided to build a pipeline around images of fluorescent DNA counterstains. While this approach does not specifically label the host cells or parasites, the amastigote nuclei and kinetoplasts are distinguishable from the host nuclei. This is generally used to quantify the images and previous automation methods have been based on these dyes. We selected the training data trying to capture the diversity inherent to these experiments, including images of varying quality and magnification, different formats and from different microscopes.

Implementing an algorithm that could replicate or improve the results obtained by traditional manual quantification presented three main challenges: identifying the individual objects in the image, distinguishing whether each object corresponds to a parasite or a host cell, and finally assigning each parasite to a single host cell.

To address the first challenge—accurately identifying individual objects in the images—we implemented Cellpose-SAM as the base segmentation tool (18). This model, originally trained on large datasets of fluorescently labeled cells, performs well with DNA counterstains and requires only minimal fine-tuning to adapt to our data. Cellpose-SAM provided a foundation for the subsequent steps of the pipeline.

To solve the second problem, we decided to adjust two separate models capable of detecting either the host nuclei or the parasites’ signal. Starting from Cellpose-SAM, we first used it to segment the nuclei of infected cell cultures with a fluorescent DNA counterstain. The base model performs well at detecting nuclei but is prone to errors when processing the amastigotes’ signal, often detecting them as separate instances or sometimes including them as part massive nuclei. This was corrected through the Cellpose GUI with a human-in-the-loop training scheme using the default parameters (Fig 1). The training consisted primarily of eliminating instances including amastigotes, while also correcting some masks for overlapped nuclei. This finally resulted in a decrease in overlap with a specific anti-*T. cruzi* stain from 8.63% to 1.81% in the validation data (S1 Fig).

**Figure 1:**
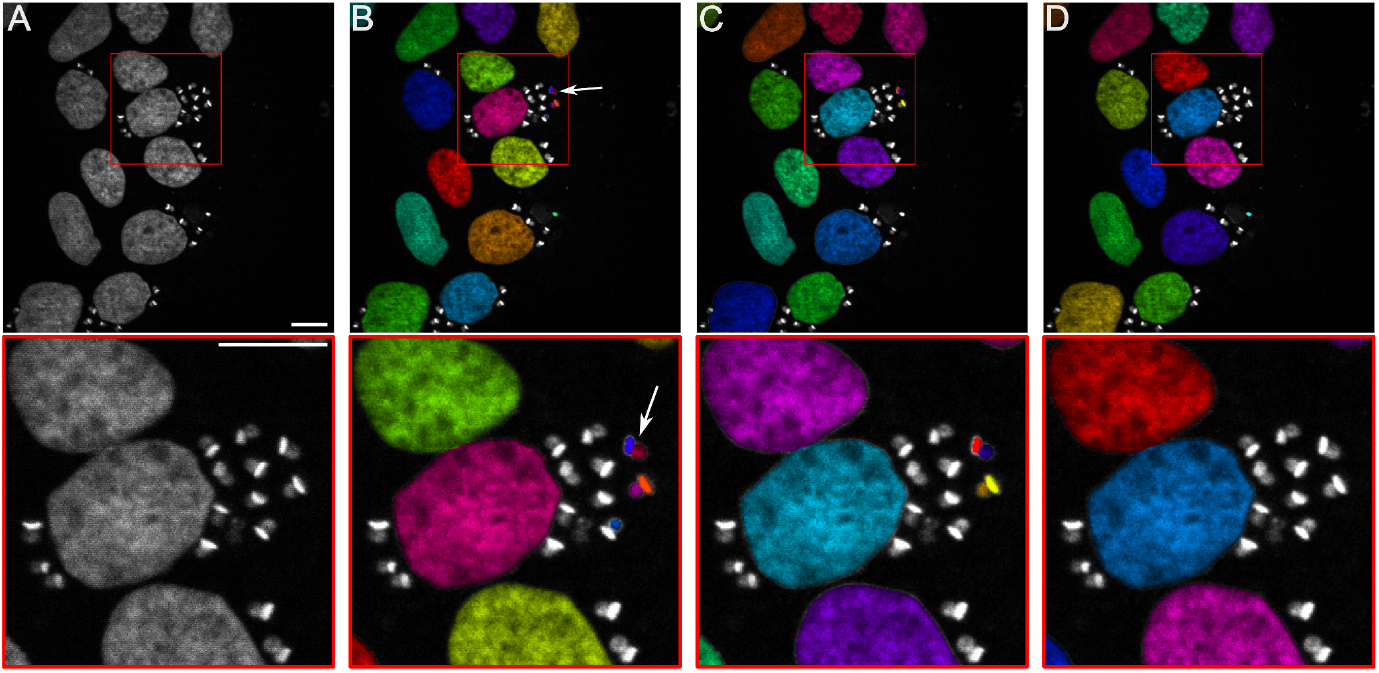
Host-nuclei model refinement. A: Grayscale image of BeWo cells infected with Dm28c parasites and stained with Hoechst 33342. B: segmentation with cellpose-sam. C: segmentation with a partially trained model. D: segmentation with the model that was used for validation. Red frames correspond to the insets. Scale bars = 20 μm. The arrows indicate an instance of inaccurate segmentation. Each colour corresponds to a unique segmentation instance.

### Model training for *T. cruzi* amastigote identification

We followed a similar approach to segment the parasites. Because the base model mostly ignores the signal corresponding to the amastigotes, the initial training consisted of eliminating the host-nuclear masks that the model generated and instead annotating the amastigotes (Figs 2A and 2B). These ROIs were drawn to include both the parasite nucleus and kinetoplast whenever both were distinguishable and, when available, a second channel with a specific anti-*T. cruzi* stain was used to confirm whether these truly were parasite structures. However, no masks were drawn where there was no evident signal in the DNA channel and the second channel was not otherwise used while fine-tuning the model.

**Figure 2:**
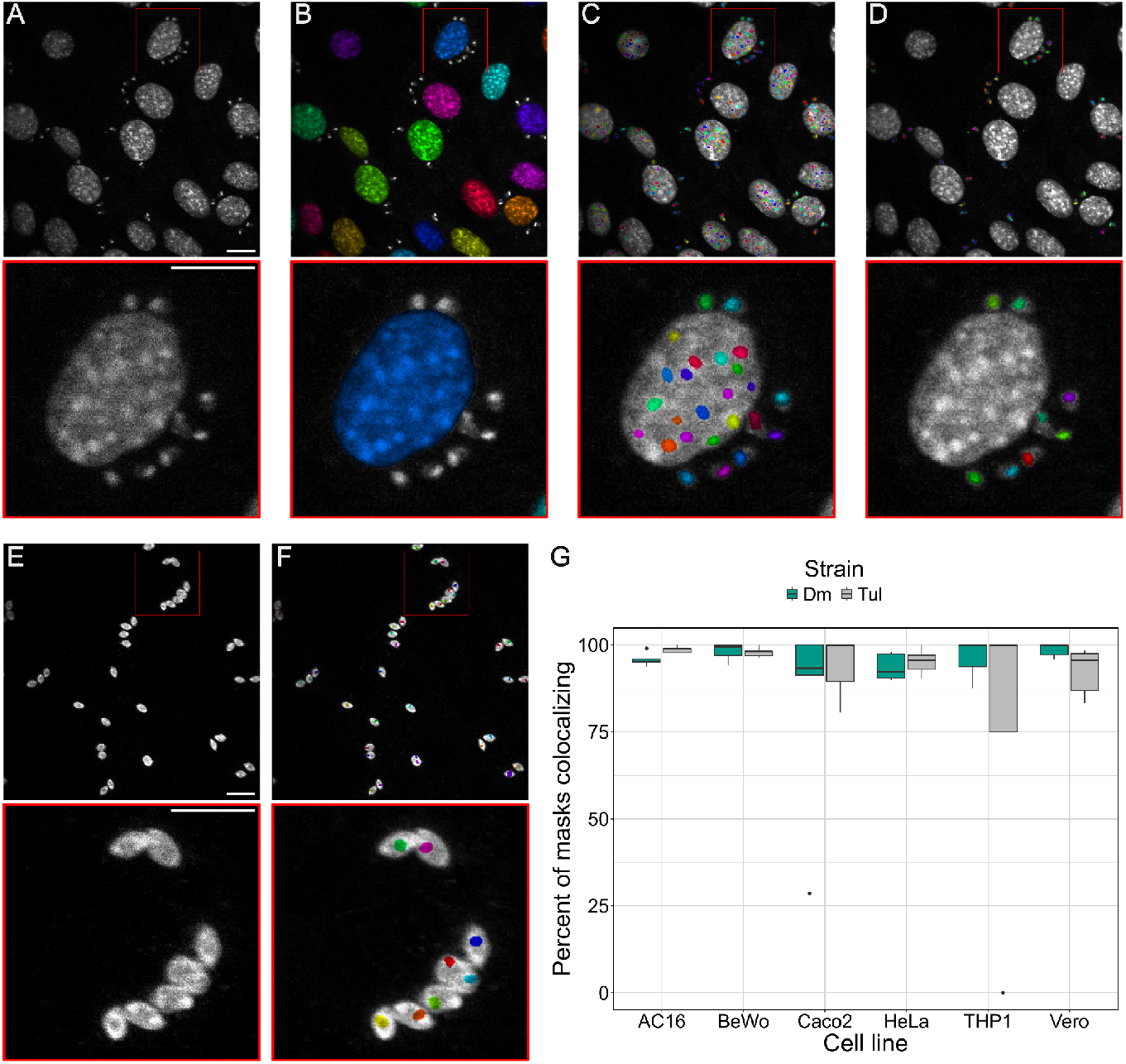
*T. cruzi* model refinement. A: grayscale image of AC16 cells infected with Dm28c parasites and stained with Hoechst 33342. B: segmentation with Cellpose-SAM. C: segmentation with a partially trained model. D: segmentation with the model that was used for validation. E: grayscale image of the specific anti-*T. cruzi* channel. F: the masks that were generated on the nuclear channel by the final model were overlaid onto the specific channel. G: the percentage of parasite masks that colocalized with the specific stain for each cell line-strain group. Red frames correspond to the insets. Scale bars = 20 μm. Each colour corresponds to a unique segmentation instance. Dm: infection with Dm28c trypomastigotes. Tul: infection with Tulahuen trypomastigotes.

After the initial round of training, our model ignored host cell nuclei while detecting smaller structures regardless of their context, with some instances being detected within the host nuclei and, in some cases, separating amastigotes into nuclei and kinetoplasts (Fig 2C). This behaviour was corrected with a second batch of training images, resulting in single masks for each pair of nucleus and kinetoplast and nuclear structures being eliminated. This generated a bias towards detecting kinetoplasts in cases where both structures were clearly distinguishable. A final round of training was used to remove false positives that seemed to detect imaging noise or staining artifacts (Fig 2D). The model was evaluated against the images in the validation dataset by analyzing the colocalization between the masks generated on the nuclear channel and the specific anti-*T. cruzi* channel (Fig 2E and 2F). On average, 93.14% of masks colocalized with the specific stain (Fig 2G). The largest error was observed with some images of THP-1 cells infected with Tulahuen parasites, which upon closer inspection seemed to be non-specifically stained on the Hoechst channel. The number of parasite objects detected on images from uninfected cells did not differ significantly compared to a manual quantification.

### Algorithm development for quantification of infection rates and parasite burden

Once both models showed a satisfactory performance on the training data, we developed an algorithm around them to quantify the infections. We first began with two rounds of instance segmentations: one using the nuclear model and another using the amastigote model. This generated two separate sets of objects for each image: the host cell nuclei and the parasites. Once all the objects of each type were detected for the whole batch of images, we proceeded with quantifying each individual image. For each parasite in the field-of-view, we calculated the distance between the amastigote’s centroid and every nuclei’s centroid in the image. Each parasite was then tagged according to its closest nucleus (Fig 3). Finally, we iterated over each nucleus and recorded the number of parasites tagged with its identifier. This approach is similar to that employed by Yazdanparast et al. (7). The results for a batch of images were recorded in two dataframes: one containing the number of amastigotes in each cell and another with the summary for each image. We also found it useful to save a visual summary of how each image was analysed, showing the masks that were generated by each model, how the parasites were assigned to the nuclei and the summary metrics for the image.

**Figure 3:**
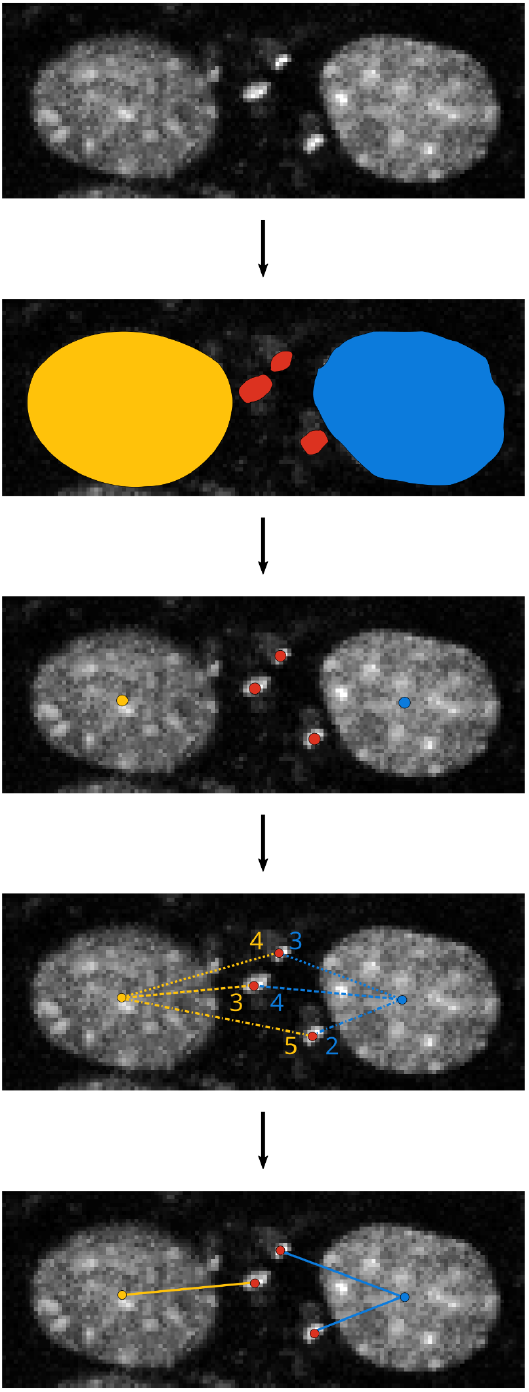
schematic of the algorithm employed to assign amastigotes to host-cells. Briefly, host-nuclei (blue and yellow) and parasites (red) were segmented separately. From these masks, the coordinates for the centroid of each object was obtained. Then, the distances between every amastigote centroid and every host-nucleus centroid were calculated. Each amastigote is then assigned to the closest host-cell.

### Algorithm validation

To validate the algorithm, we used a set of confocal images from 6 cell lines infected with two different parasite strains, showing various staining characteristics and nuclear morphologies. The algorithm’s overall performance was evaluated against a manual quantification of this dataset. We first compared the total number of objects detected in each image by calculating the symmetric mean absolute percentage error (SMAPE), for amastigotes and host nuclei separately. This metric allowed us to gauge the difference between both methods, relative to the average number of objects in each image.

Fig 4A shows the relative error in the number of parasites for each image. In general, the amastigote model tends to somewhat underestimate the number of parasites compared to the manual quantification. However, this error seems to be small in most cases and the median quantification for most groups is around 5% of the manual count. Outside the general tendencies, there are two stand-out groups: Caco2 cells infected with Dm28c parasites and THP-1 cells infected with Tulahen parasites, the same groups that previously underperformed when compared against the anti-*T. cruzi* stain. The underestimation in the latter group can be attributed to the previously mentioned staining problems. As for the former group, some images with very few objects and low signal-to-noise ratio resulted in 6 false positives that translated to a large relative overestimation. However, errors caused by few objects can be diluted in the overall quantification. Interestingly, on a smaller dataset of HeLa cells, the total number of amastigotes diverged more between two human operators than between the operator that trained the model and the model itself (differences of -14% and 6%, respectively). The host-nucleus model gave extremely similar results with both counting methods (Fig 4B).

**Figure 4:**
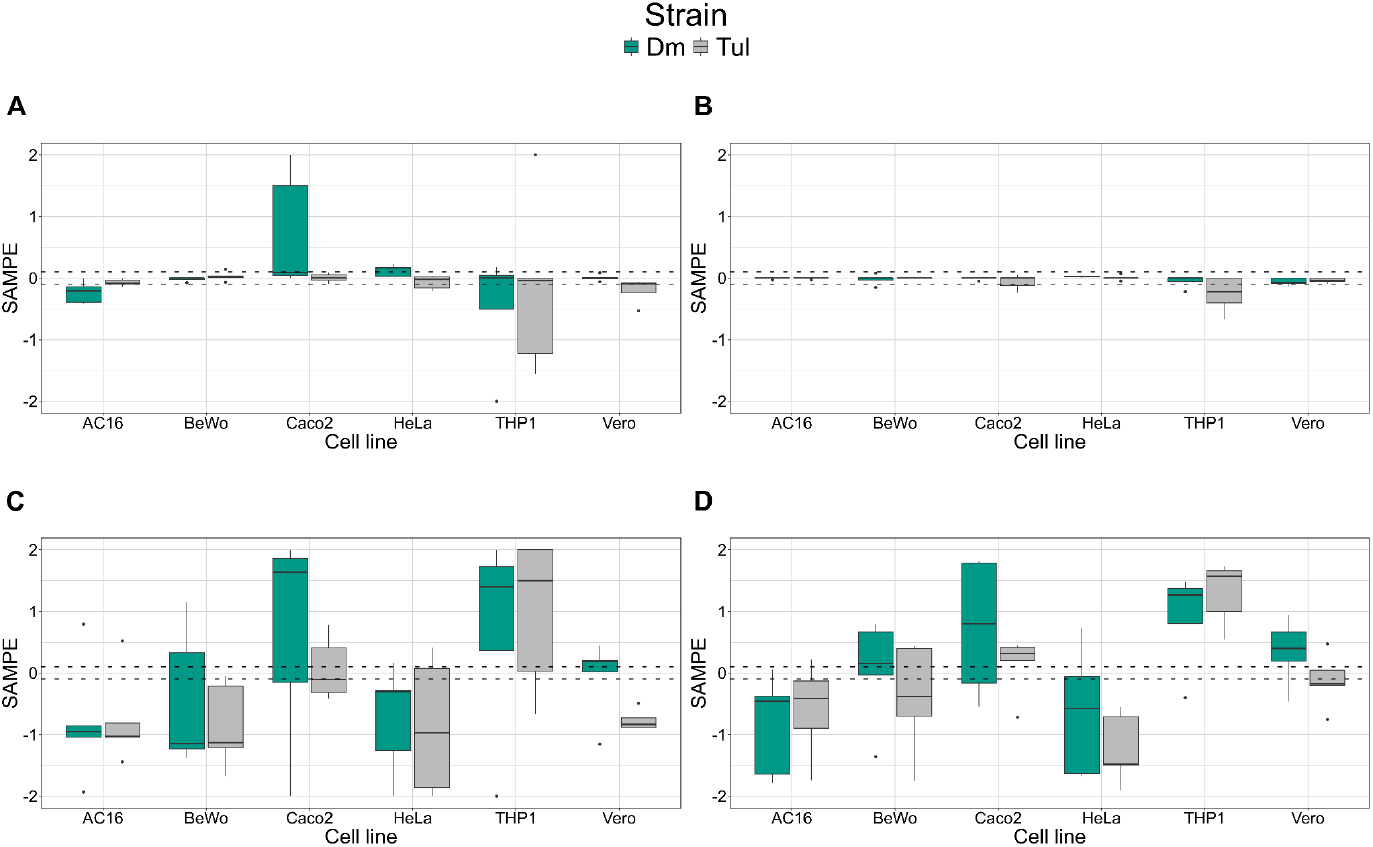
Symmetric Mean Absolute Percentage Error (SMAPE) of automated quantification algorithms compared to the manual quantification of the validation dataset, calculated for each individual image of the Cell-Strain groups. A: NN model for amastigote detection. B: NN model for host-nuclei detection. C: morphological image analysis method for amastigote detection. E: morphological method for host-nuclei detection. Dm: Dm28c infection. Tul: Tulahuen infection. The dashed lines around 0 bound 5% deviations.

To understand how the models compare to more traditional morphological image analysis techniques, we replicated the algorithm designed by Yasdanparast et al., to the best of our abilities. Overall, there seem to be much larger deviations from the manual quantification using this methodology, for both amastigotes and host cell nuclei (Figs 4C and 4D, respectively). The deviations seem to depend more on the cell line, suggesting that each one needs specific techniques and parameters to be appropriately analyzed. In fact, the error for amastigote detection is strongly correlated with the error for host-nuclei detection (0.72 Pearson coefficient, compared to 0.30 using the NN-based methodology). This is in accordance with nuclei detection being necessary for amastigote detection with the morphological methodology, and highlights the benefit of independently segmenting for both types of objects. It should be noted that, while quantifications with the morphological method deviate from manual counts, the overall conclusions from the analyses may be the same.

The quantification of the test images is shown in Fig 5, comparing the automated quantification to the manual quantification. The main measure that was obtained was the number of amastigotes in each cell, which is summarized in Fig 5A. Overall, the mean number of amastigotes per cell was similar between methods for all cell line and parasite strain pairs, which suggests that errors that were previously observed became diluted as the data from all fields was accumulated; the detection of both host nuclei and parasites by the automated method seems to be comparable to manual counting. The distribution of the data was also similar between methods, with most groups showing a larger density of cells with little to no parasite burden, with a decreasing density of more infected cells, although there are some deviations.

**Figure 5:**
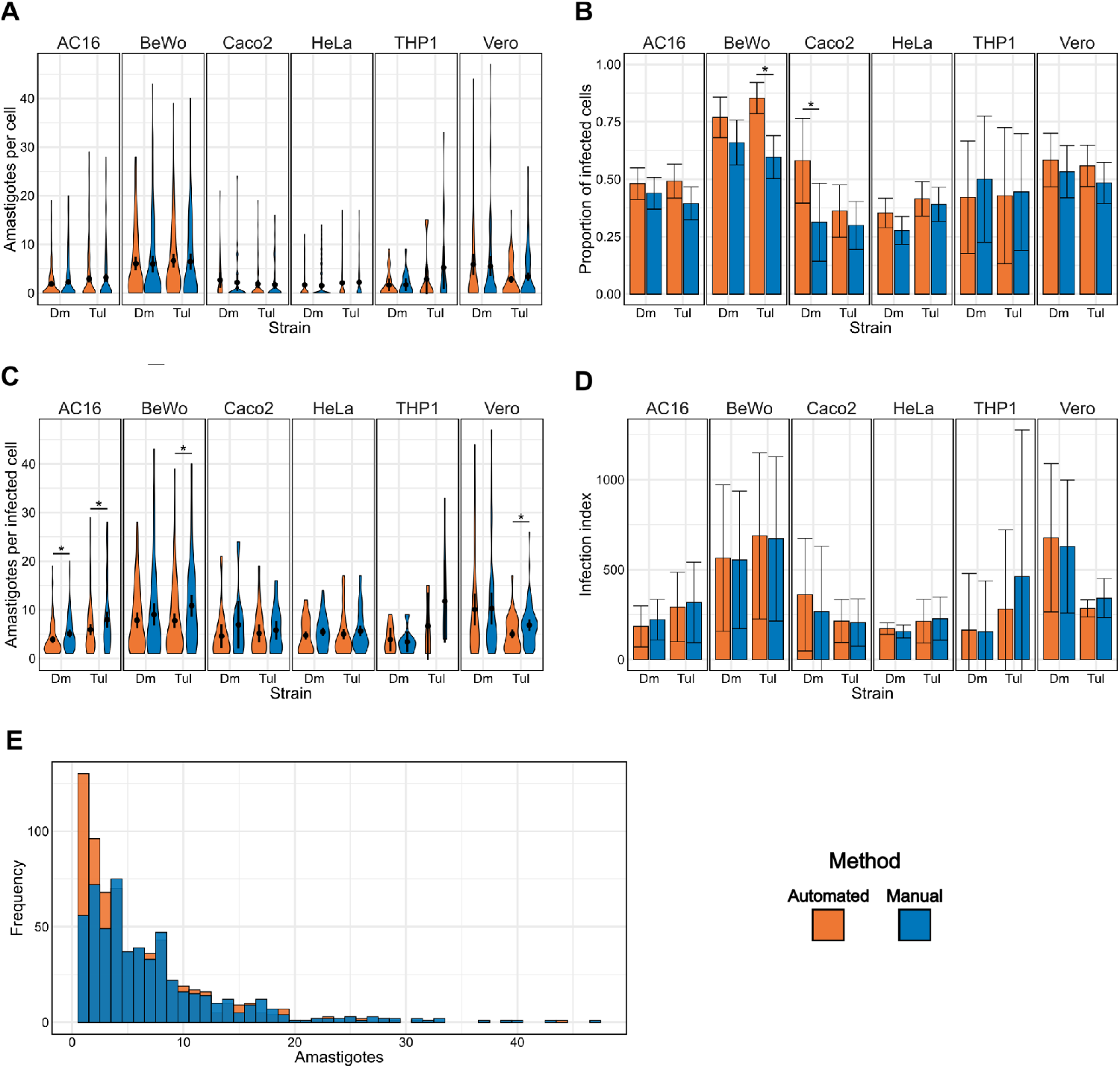
Quantification of the validation dataset using the automated (NN-based) and manual methods, grouped by cell line and the parasite strain used to infect (Dm: Dm28c; Tul: Tulahuen) . A: violin plots of the number of amastigotes in each individual host-cell. B: average proportion of infected cells in each image. C: the number of amastigotes in infected cells. D: average infection index calculated for each image. E: total histogram of the number of amastigotes in infected cells. The total number of amastigotes counted was 3927 and 4128 for the automated and manual methods, respectively. Asterisks represent contrasts with a p-value < 0.05.

There seems to be a tendency for the algorithm to overestimate the proportion of infected cells relative to the manual quantification (Fig 5B). This tendency was exaggerated for the BeWo cells infected with Tulahuen parasites and for the Caco2 cells infected with Dm28c parasites. The overall effect could be attributed to the assignment criterion being based on distance alone, with no regards for how the parasites cluster together inside their host cells. The BeWo and Caco2 lines are fairly large cells that can grow in tight clusters where it is difficult to distinguish the limits between the cytoplasms of different cells, so it stands to reason that highly infected cultures register some parasites as closer to a different cell than its true host. This overdispersion can also be observed in the number of parasites per infected cell (Fig 5C). Here, there is a general tendency towards underestimation, consistent with the overestimation of the total number of infected cells and further confirming that distance assignment seems to distribute the parasites more uniformly over all cells. The infection indexes do not seem to be affected by the difference in assignment, although some valuable information might be lost (Fig 5D).

The differences in assignment criteria between the algorithm and the operator can be better observed in the overall histogram for amastigotes per infected cells (Fig 5E). Here, the automated method accumulates over twice the amount of cells that are infected with a single parasite, compared to the manual quantification. Again, this suggests a more uniform dispersion of the amastigotes amongst the detected cells caused by the distance assignment. The frequency for the automated method then decays following the expected negative binomial distribution, while the manual method shows a flatter, multi-modal data distribution highlighting the operator’s particular assignment criteria that does not assume independence between the amastigotes. This highlights the chief advantage of manual quantification: a human operator is more likely to understand how the amastigotes are clustered in an infection and assign them more accurately, although this also introduces a certain bias to the analysis. While we have here highlighted the differences between the methods, the divergence between the results is rather small in most cases, making it difficult to ascertain which method is generally more accurate for quantifying our validation dataset using only DNA counterstains.

## Discussion

In this study, we developed and validated a neural network (NN)-based algorithm for the automated quantification of intracellular *Trypanosoma cruzi* infections in mammalian cells using simple DNA counterstains. This approach directly addresses a critical methodological gap in Chagas disease research, where infection quantification is still largely dependent on manual counting or on methods that require genetically modified parasites. By combining two independently trained models—one for host cell nuclei and another for amastigotes—and integrating them into a unified pipeline, we provide a tool capable of delivering objective, reproducible, and high-throughput measurements of infection rates and parasite load. Such advances are essential for strengthening early-stage drug discovery pipelines and reducing reliance on labor-intensive manual quantification.

The adoption of AI-based analysis has proven successful in other parasitic diseases, such as malaria and leishmaniasis, where cellular phenotypes and subcellular localization patterns are critical (19,20). For instance, Portella and co-workers (9) described the use of automated microscopy and analysis pipelines for assessing infection in stem-cell-derived cardiomyocytes, demonstrating that such models can simulate physiologically relevant host environments while being compatible with scalable image-based quantification. In the context of Chagas disease, leveraging CNN-based tools for automated quantification of intracellular *T. cruzi* offers a scalable and objective solution for early-stage drug screening pipelines. Such platforms allow for the integration of multi-parametric analyses, including infection indexes, parasite replication rates, and host cell viability, providing a comprehensive assessment of drug activity. Moreover, standardization enabled by algorithm-driven image analysis reduces operator bias and facilitates inter-laboratory comparisons, which are essential for validating compound efficacy and advancing lead candidates through the drug discovery pipeline.

Coupling our NN-based pipeline with high-throughput imaging (HTI) systems could further expand its utility. HTI platforms integrate automated microscopy with large-scale image acquisition, enabling rapid screening of compound libraries while maintaining single-cell resolution. When combined with AI-driven analysis, these systems have proven effective in parasitic disease drug discovery and phenotypic screening, offering both scalability and reproducibility across diverse experimental conditions (21,22). Incorporating HTI into our workflow would allow for accelerated evaluation of compound efficacy, facilitate multi-parametric readouts, and strengthen the translational potential of image-based assays for Chagas disease.

The algorithm showed robust performance when compared to manual quantification, which remains the most common approach. Overall, the differences between both methods were small, with median deviations around 5% of manual counts, and the distribution of parasite burden per cell was similar across datasets. Importantly, the independent segmentation of host nuclei and parasites reduced error propagation and improved reproducibility compared to morphological pipelines such as INsPECT (7). While INsPECT reached similar conclusions, it tended to inflate absolute values and required more complex parameter adjustments. In contrast, our NN-based method was easier to implement, less sensitive to acquisition parameters, and provided consistent results across different cell lines and parasite strains. Moreover, the original INsPECT software is no longer readily available through the published link, requiring us to build a version of the program following the author’s description.

An important aspect of the NN-based pipeline was the choice of segmentation tool. There are currently several pretrained models designed to detect objects in microscopy images from biological samples. We chose Cellpose as it is specifically designed to segment fluorescent images of cells with little need for parameter adjustment, using vector gradients instead of mask predictions which allows for better accuracy in the final segmentation. Cellpose was also originally trained on a large dataset which included fluorescently labeled cells, including fluorescent DNA stains (17). The available Graphical User Interface (GUI) allows for fine-tuning existing models on the user’s data, without the need for specialized knowledge (23). While the original model was based on the U-Net architecture, a convolutional autoencoder, it has now adopted a transformer foundation model as its base (Cellpose-SAM), which greatly increases its inductive capabilities and potentially allows for diverse inputs with no parameter modifications (18). The pre-trained model was effective at detecting host nuclei but initially struggled with amastigote signals, often merging them into large nuclear masks or splitting them into fragments. These issues were corrected through additional training with curated datasets. This highlights both the adaptability of the algorithm and the importance of fine-tuning pretrained models to the specific characteristics of *T. cruzi* infections. In this sense, the pipeline benefits from the strengths of foundation models while remaining flexible enough to address the unique challenges posed by intracellular parasites.

Despite these strengths, certain limitations must be acknowledged. The algorithm showed reduced accuracy in cases of suboptimal staining, particularly in THP-1 cells infected with Tulahuen parasites, and in images with low signal-to-noise ratios or very few detectable objects. These edge cases emphasize the importance of careful dataset curation and suggest that further refinement of training data, including broader representation of staining conditions and cell types, could improve generalizability. Additionally, the algorithm’s tendency to distribute amastigotes more evenly across host cells than manual quantification may lead to slight overestimation of infection rates in specific contexts. Although an experienced operator is capable of adapting to such a setback, these sorts of images should be excluded from standardized quantifications inputs to maintain accuracy.

Taken together, our work shows that NN-based automated quantification of intracellular *T. cruzi* infections offers a practical and accessible solution. By achieving accuracy comparable to manual methods while offering scalability, reproducibility, and reduced operator bias, the algorithm surpasses the limitations of current approaches. Its integration into drug discovery pipelines has the potential to reduce compound evaluation timelines, harmonize protocols across laboratories, and foster collaborative efforts aimed at identifying novel therapeutics for this persistent and underserved public health threat.

## Acknowledgements

S.C.V.L., and G.D.A are members of the Research Career of CONICET. J.I. is carrying out his PhD thesis at Departamento de Fisiología, Biología Molecular y Celular, Facultad de Ciencias Exactas y Naturales, Universidad de Buenos Aires.

## Supporting information

**Figure S1:**
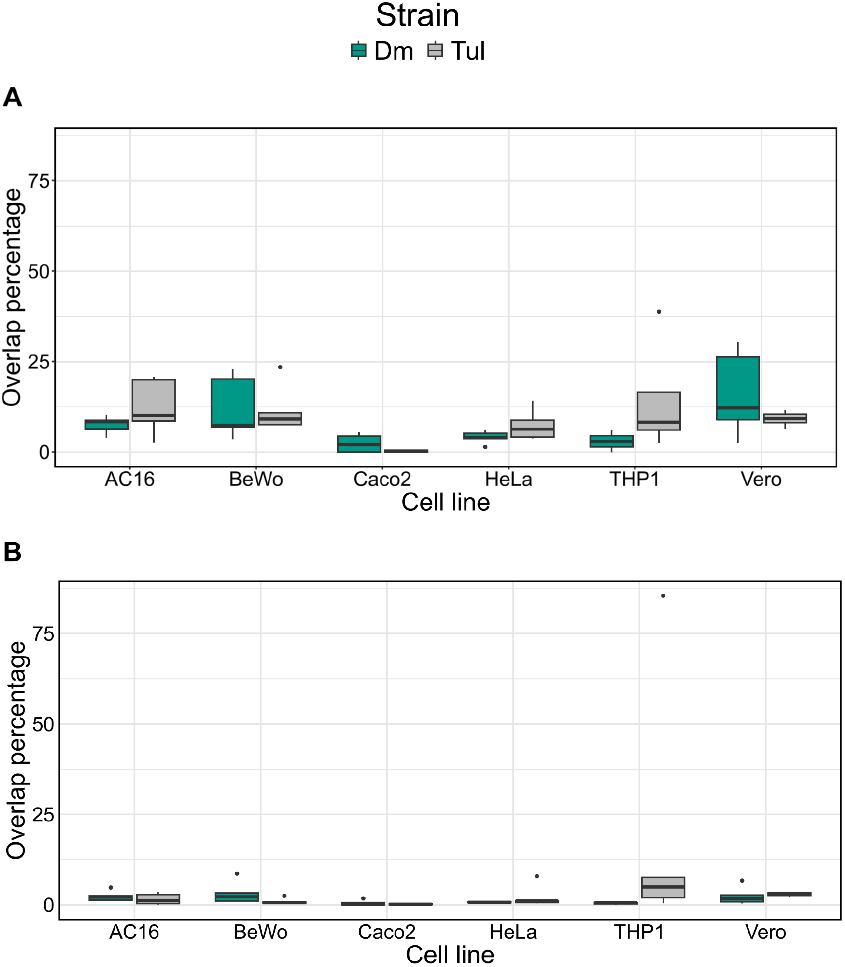
The percentage of the total surface area of the host-nuclei masks that overlapped with the specific signal form the anti-*T. cruzi* serum, for each Cell line-Strain group. A: masks were generated with the Cellpose-SAM model. B: masks were generated with the refined model. Dm: infection with Dm28c trypomastigotes. Tul: infection with Tulahuen trypomastigotes.

